# Identification of a genetic region in linked to tolerance to MRSA infection using Collaborative Cross mice

**DOI:** 10.1101/2023.07.13.548933

**Authors:** Aravindh Nagarajan, Kristin Scoggin, L. Garry Adams, David Threadgill, Helene Andrews-Polymenis

**Affiliations:** Interdisciplinary Program in Genetics and Genomics, Texas A&M University, College Station, Texas, United States of America; Department of Molecular and Cellular Medicine, Texas A&M University, College Station, Texas, United States of America; Department of Microbial Pathogenesis and Immunology, Texas A&M University, College Station, Texas, United States of America; Texas A&M Institute for Genome Sciences and Society, Texas A&M University, College Station, Texas, United States of America; Department of Biochemistry & Biophysics and Department of Nutrition, Texas A&M University, College Station, Texas, United States of America; Department of Veterinary Pathobiology, Texas A&M University, College Station, Texas, United States of America

**Keywords:** Host-pathogen interaction, MRSA USA300, tolerance, QTL mapping, F2-Cross, *C5a*-*C5ar* axis.

## Abstract

*Staphylococcus aureus* (*S. aureus*) colonizes humans asymptomatically but can also cause opportunistic infections, ranging from mild skin infections to severe life-threatening conditions. Resistance and tolerance are two ways a host can survive an infection. Resistance is limiting the pathogen burden, while tolerance is limiting the health impact of a given pathogen burden. In previous work, we established that collaborative cross (CC) mouse line CC061 is highly susceptible to Methicillin-resistant *S. aureus* infection (MRSA, USA300), while CC024 is tolerant. To identify host genes involved in tolerance after *S. aureus* infection, we crossed CC061 mice and CC024 mice to generate F1 and F2 populations. Survival after MRSA infection in the F1 and F2 generations was 65% and 45% and followed a complex dominant-recessive inheritance pattern. Colonization in F2 animals was more extreme than in their parents, suggesting successful segregation of genetic factors. We identified a QTL peak on chromosome 7 for survival and weight change after infection. In this QTL, the WSB allele was present in CC024 mice and contributed to their MRSA tolerant phenotype. Two genes, *C5ar1* and *C5ar2*, have high-impact variants in this region. The complement factor, *C5a*, is an anaphylatoxin that can trigger a massive immune response by binding to its receptors, *C5ar1* and *C5ar2*. We hypothesize that *C5a* may have altered binding to variant receptors in CC024 mice, reducing damage caused by the cytokine storm and resulting in the ability to tolerate a higher pathogen burden and longer survival.

**Importance:** *Staphylococcus aureus* causes a wide range of diseases in humans. Resistance and tolerance are two ways a host can survive an infection. Resistance is limiting the pathogen burden, while tolerance is limiting the health impact of a given pathogen burden. Tolerance mechanisms are poorly understood in context of host-pathogen interaction. To identify host genes involved in tolerance after *S. aureus* infection, we crossed CC061 mice and CC024 mice. The genetic factors controlling tolerance were well segregated in the F2 population. Using QTL mapping, we identified a significant peak on chromosome 7 for survival and weight change after infection. Two genes, C5ar1 and C5ar2, have high-impact variants in this region. We hypothesize that C5a may have altered binding to variant receptors in CC024 mice, reducing damage caused by the cytokine storm and resulting in the ability to tolerate a higher pathogen burden and longer survival.

## Introduction

*Staphylococcus aureus (S. aureus*) is a gram-positive coccus well adapted to humans and a variety of companion, farm, and wild animals (Mrochen, Fernandes de Oliveira, Raafat, & Holtfreter, 2020). *S. aureus* is a commensal organism as well as an opportunist that can cause severe morbidity and mortality (Krismer, Weidenmaier, Zipperer, & Peschel, 2017; Lowy, 1998). Globally, *S. aureus* is one of the leading causes of bacterial-related hospitalizations and is responsible for more than one million deaths annually (GBD, 2022). *S. aureus* is a growing cause of concern because it spreads easily, and many isolates are resistant to several classes of antibiotics (Otto, 2013; Turner et al., 2019).

Nasal carriage of *S. aureus* varies widely across the human population (Eriksen, Espersen, Rosdahl, & Jensen, 1995; Mehraj et al., 2016). Not all colonized individuals develop infections, but colonization is a risk factor for skin and soft tissue infections (Clarridge, Harrington, Roberts, Soge, & Maquelin, 2013; Howden et al., 2023). These superficial infections are precursors for 50% of *S. aureus* bacteremia (SAB) (Yarovoy, Monte, Knepper, & Young, 2019), and approximately one-third of SAB cases lead to sepsis (Fowler et al., 2003; Ringberg, Thorén, & Lilja, 2000). This variability in disease outcomes after *S. aureus* colonization, suggests that this host-pathogen interaction may depend on both bacterial and host genetics.

Resistance and tolerance are two ways a host can survive an infection. Resistance is defined as maintaining health by limiting the pathogen burden, while tolerance is limiting the health impact of infection despite a high pathogen burden (Schneider & Ayres, 2008). Mechanisms underlying tolerance to infection have recently been described for parasitic (Gozzelino et al., 2012; Seixas et al., 2009) and viral infections (Irving, Ahn, Goh, Anderson, & Wang, 2021). However, mechanisms underlying tolerance to bacterial infections are poorly understood (Barman & Metzger, 2021).

The CC is a large panel of genetically diverse recombinant mouse strains derived from eight founder strains (five classical inbred strains - A/J, C57BL/6J, 129S1/SvlmJ, NOD/ShiLtJ, NZO/HlLtJ, and three wild-derived strains - CAST/EiJ, PWK/PhJ, and WSB/EiJ). This collection captures approximately 90% of the genetic diversity in laboratory mice (Churchill et al., 2004; Roberts, Pardo-Manuel de Villena, Wang, McMillan, & Threadgill, 2007; Threadgill, Miller, Churchill, & de Villena, 2011). With fully sequenced genomes, CC strains are ideal candidates for Quantitative Trait Locus (QTL) mapping (Shorter et al., 2019; Srivastava et al., 2017). Crosses between CC strains have been successful in identifying genetic loci underlying susceptibility to seizure (Gu et al., 2020), and *Mycobacterium tuberculosis* (Smith et al., 2022), and *Salmonella* Typhimurium (J. Zhang et al., 2019) infections.

Host genetic variation plays role in sensitivity, tolerance and resistance mechanisms (Albers et al., 1987; Wambua et al., 2006) (Hill, 1998; Malo & Skamene, 1994). We created a large F2 intercross panel by crossing collaborative cross (CC) strains with known phenotypes with respect to MRSA USA300 infection, using a MRSA susceptible strain (CC061) and a tolerant strain (CC024). In our previous work we determined that parental CC061 mice rapidly lost weight and met our euthanasia criteria, while CC024 mice lost less weight and survived the study period. Cytokine and gene expression patterns differed between these strains both before and after infection.

Survival in F1 and F2 generation followed a complex dominant-recessive pattern of inheritance. Colonization in F2 animals was more extreme than in the parental strains, suggesting successful segregation of genetic factors. We identified a QTL peak on chromosome 7 for survival and weight change after infection. The genes within the QTL region suggest a mechanism for tolerance to MRSA infection.

## Results

### CC061 and CC024 differ in survival after infection with MRSA USA300

In our previous study, two CC strains, CC061 and CC024, were infected with MRSA USA300 and monitored for a week after infection (Submitted: mBio01812-23). All of the CC061 mice met our euthanasia criteria with a median survival of three days and were classified as susceptible (Fig 1A). All the CC024 mice survived until day 7 with high organ colonization and were classified as tolerant (Fig 1A and 1B). CC061 mice lost 18% of their body weight on average (Fig 1C). Weight loss in CC024 mice was variable, three animals gained weight and three lost weight, a mean of -2.5% of body weight was lost across CC024 mice (Fig 1C). There was no sex difference in survival or weight loss in these strains, and the kidney was the most heavily colonized organ in both CC024 and CC061 mice. Kidney, heart, and lung colonization did not significantly differ between CC061 and CC024 (Fig 1B). Spleen and liver colonization were higher in CC061 than in CC024 mice, yet there was no difference in tissue damage scores in these two strains for the spleen and liver (Fig 1D). Paradoxically, we observed more severe damage in the kidney in tolerant CC024 mice than in sensitive CC061 mice (Fig 1D). This finding contradicts the previous dogma that the ability to reduce tissue damage may be a feature of the tolerance phenotype (Medzhitov, Schneider, & Soares, 2012; Wu & Reddy, 2017).

**Fig 1:**
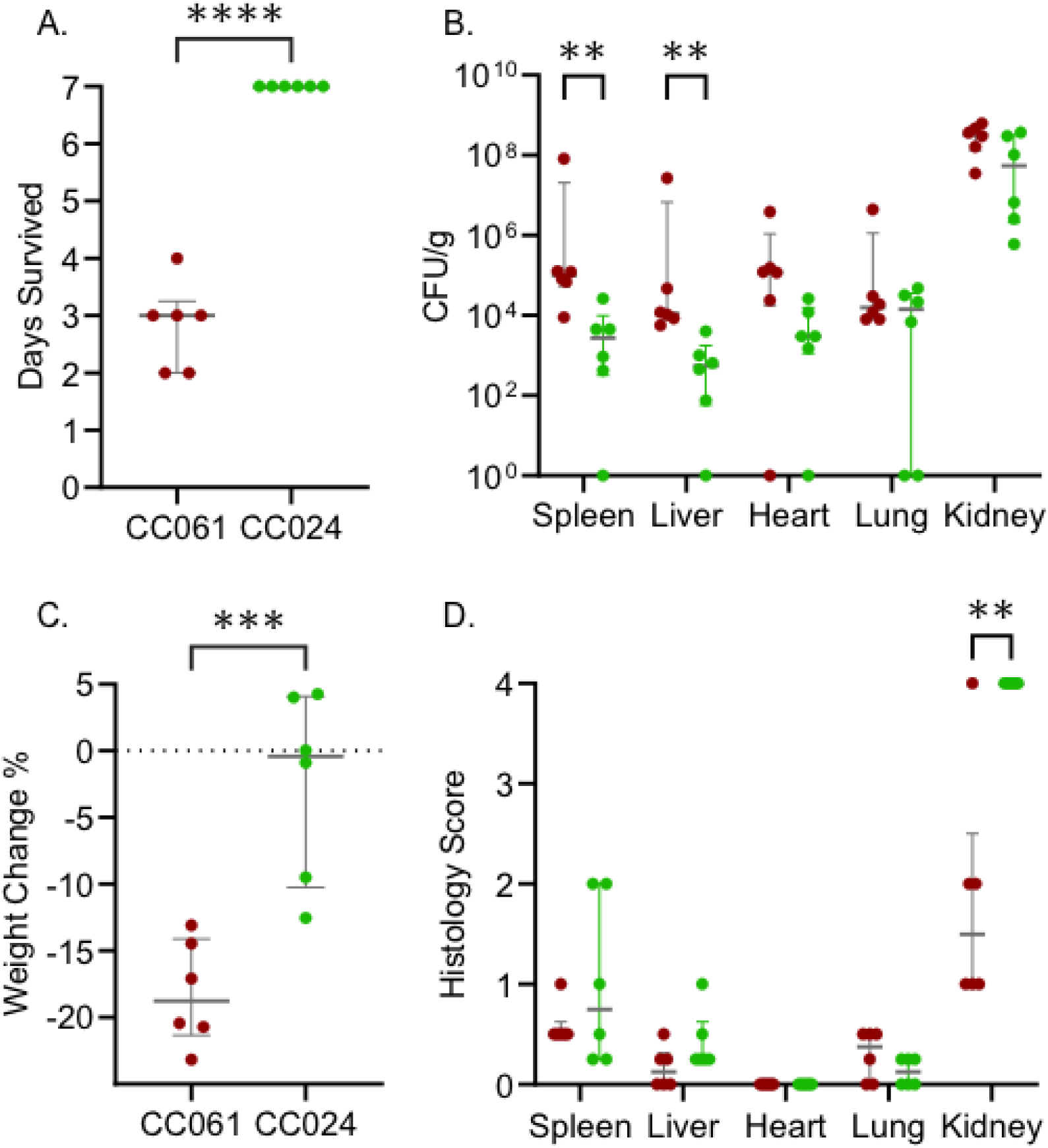
CC061 and CC024 differ in survival after infection with MRSA USA300. After intravenous infection with MRSA USA300 as described in our previous work, we show: A. Survival time B. Percent weight change after infection C. Colonization and D. Tissue damage in the spleen, liver, heart, lung, and kidney. Dots represent individual mice; red dots represent CC061; green dots represent CC024. The median and interquartile range are shown for each strain. A, B, and D – Student’s t-test and C – Mann-Whitney tests were performed (* = P < 0.05, ** = P < 0.01, *** = P < 0.001, **** = P < 0.0001)

### Immune profile differences between CC061 and CC024

Each CC strain has a unique immune profile (Collin, Balmer, Morahan, & Lesage, 2019; Graham et al., 2017), so we examined the differences in immune parameters between CC024 and CC061. CC024 mice have higher baseline circulating white blood cells (WBC) before infection than CC061 mice, driven by higher numbers of lymphocytes (LYM) (Fig 2A). After infection, both WBC and LYM were elevated in CC024 mice compared to CC061 mice (Fig 2A), but the change from baseline levels was similar in both lines (Fig 2B).

**Fig 2:**
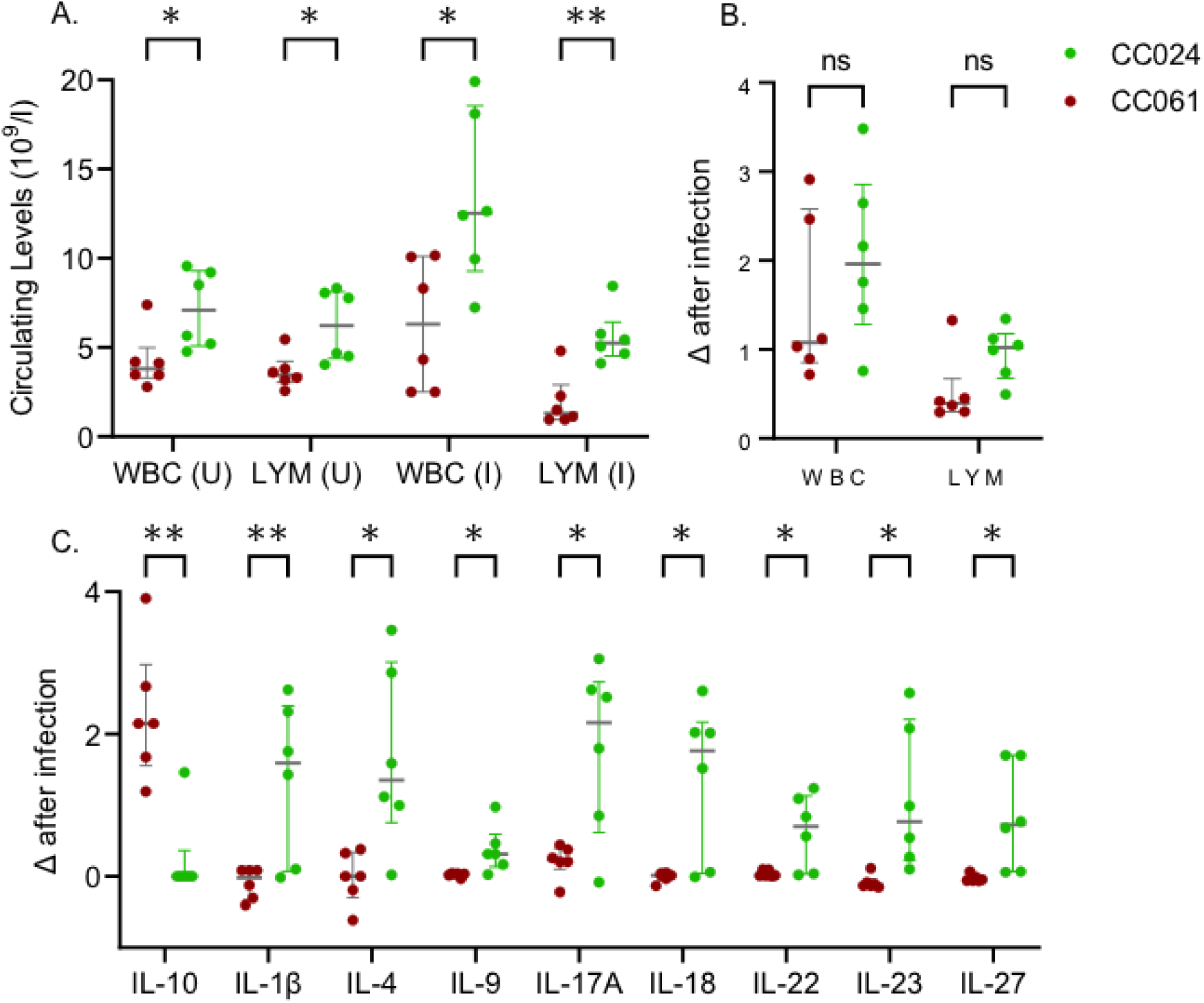
Immune profile differences between CC061 and CC024. A. Circulating levels of White Blood Cells (WBC) and Lymphocytes (LYM) in uninfected (U) and infected (I) mice. B. Change in WBC and LYM levels post-infection. C. Change in chemokine levels post-infection. Dots represent individual mice; red dots represent CC061; green dots represent CC024. The median and interquartile range are shown for each strain. Student’s t-test was performed to test for significance (ns – no significance, * = P < 0.05, ** = P < 0.01)

After accounting for strain differences in baseline levels of these cytokines, nine cytokines differ significantly between CC024 mice and CC061 mice post-infection (Fig 2C). IL-10 was significantly increased in CC061 mice after MRSA infection, while IL-1β, IL-4, IL-9, IL-17A, IL-18, IL-22, IL-23, and IL-27 were significantly increased in CC024 mice after infection (Fig 2C). These differences in immune parameters suggest that the two strains differ in their immune response after infection.

### Gene expression differences between CC061 and CC024

We sequenced total RNA from kidneys (three males and three females) from CC061 and CC024 mice after infection to identify differences in gene expression. We also sequenced total RNA from age and sex-matched uninfected CC061s and CC024s to identify baseline differences. We compared differentially expressed genes between the two strains and looked for enriched KEGG pathways (KEGG-Kyoto Encyclopedia of Genes and Genomes). Prior to infection, the complement and coagulation cascade was the only pathway up-regulated in CC024 mice compared to CC061 mice (Fig 3A).

**Fig 3:**
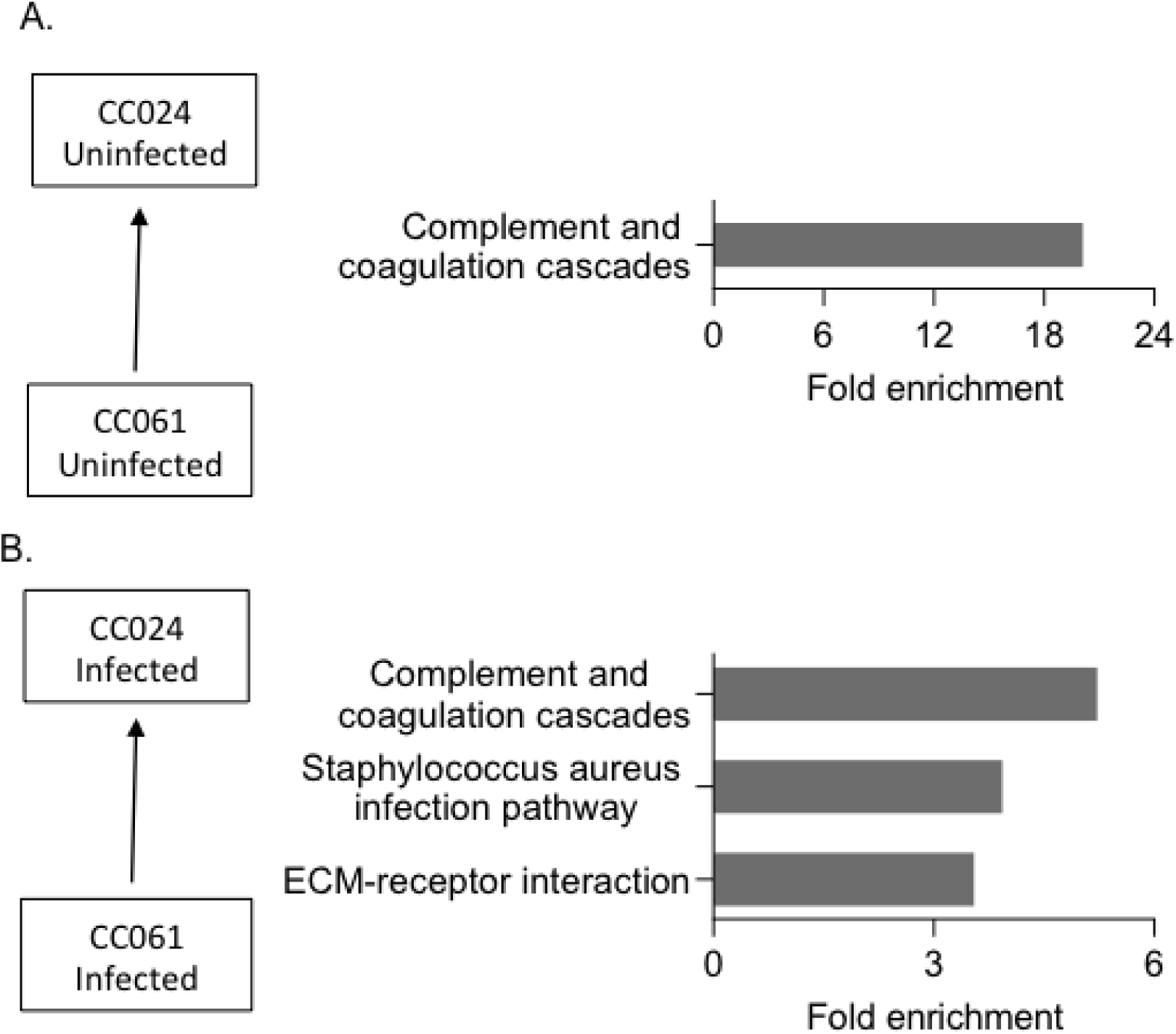
Gene expression differences between CC061 and CC024. Enriched pathways in CC024 compared to CC061 A. Before infection B. After infection. Fold enrichment was calculated by using all expressed genes as background.

Complement genes, *C3* and *C4b*, and serine/cysteine peptidase inhibitors 1a and 1d contributed to this 20-fold enrichment. The complement system plays an important role in protecting the host against bacterial infections, including *S. aureus* (Schifferli, Ng, & Peters, 1986; Wójcik-Bojek, Różalska, & Sadowska, 2022). Up-regulation of the complement cascade before and after infection in CC024 may play an important role in survival of CC024 mice after infection.

Three pathways were up regulated in CC024 mice compared to CC061 mice after MRSA infection (Fig 3B). In KEGG, the *S. aureus* infection pathway is part of the complement and coagulation cascade. Some notable genes that enriched these two pathways include complement components – *C1qa, C1qb, C1qc,* and *Cfh*, coagulation components – *Fgb, F2, F12, and Plg*, and serine/cysteine peptidase inhibitors *Serpina (1a-e and 2)* (Table S4). The third pathway, extracellular matrix (ECM), contained genes, Fibronectin (*Fn1*), Vitronectin (*Vtn*), and collagens – *Col1a1, Col1a2*, and *Col6a3*.

### Survival is linked to dominant alleles in CC024 mice

To understand the genetic components involved in tolerance after infection, we created an F1 population by crossing MRSA susceptible CC061 mice with MRSA tolerant CC024 mice. We infected 14 animals from the F1 generation and compared their phenotypes with those of their parents. The median survival for the F1 generation was seven days, similar to CC024 mice and significantly higher than CC061 mice (Fig 4A). The F1 mice maintained their weight similar to the CC024 parent and lost substantially less weight than the CC061 parent (Fig 4B). These data support a complex dominant-recessive genetic inheritance pattern for survival in the F1 cross between CC061 and CC024 mice. Survival after MRSA infection has a genetic component and is likely linked to dominant alleles in CC024.

**Fig 4.**
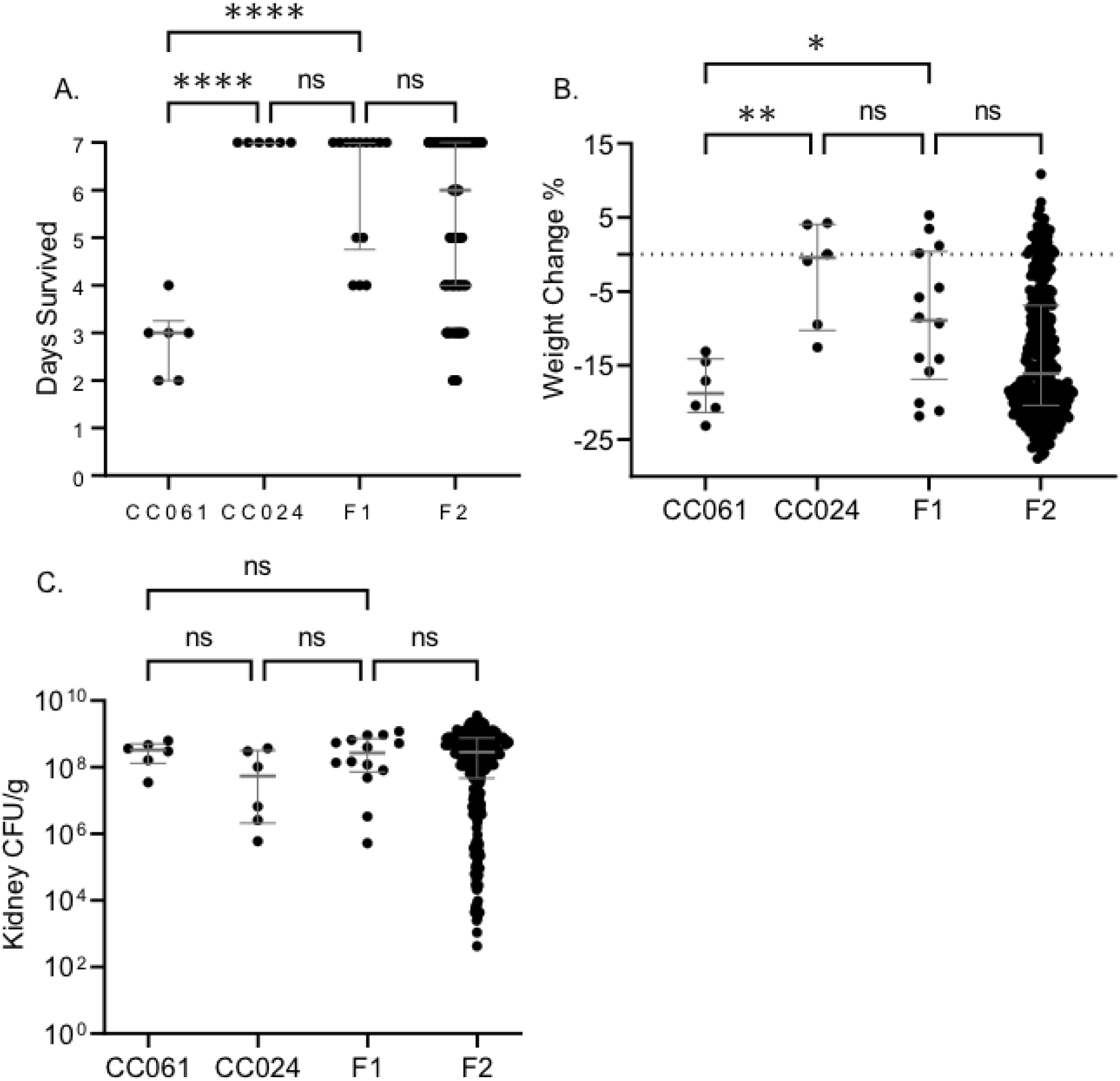
The F2 population is similar to F1 in survival and weight change after infection. A. Survival time B. Percent weight change after infection C. Kidney colonization. Dots represent individual mice. Significance tests were performed between the two parents and F1 and between F1 and F2. The median and interquartile range are shown. A and B – Student’s t-test and C – Mann-Whitney tests were performed (ns – no significance, * = P < 0.05, ** = P < 0.01, *** = P < 0.001, **** = P < 0.0001)

### F2 population is similar to the F1 generation in survival and weight change after MRSA infection

F1 mice were bred together to generate a large panel of F2 mice to further investigate the genetic components of tolerance and susceptibility. We infected 316 F2 mice with MRSA USA300. Among the 316 mice, 175 (55%) survived the 7-day study period, while the other 141 (45%) met our euthanasia criteria prior to day 7. Pre-infection weight (R = 0.04) and infection dose (R = 0.05) did not correlate with survival (Fig S1A). Weight change after infection (R =0.67) and kidney colonization (R = -0.57) correlated significantly with survival (Fig S1A). Survival in the F2 population was not influenced by sex (Fig S1B). The median survival for the F2 population was six days, very similar to the median survival of the F1 population (Fig 4A). Furthermore, the F1 and F2 generations were identical in weight change post-infection (Fig 4B), and kidney colonization was also similar between the parents, F1 and F2 generations (Fig 4C). The similarity in infection outcome between the F1 and F2 indicates that the longer survival phenotype of the tolerant CC024 strain is dominant.

### Genotype and QTL analysis confirmation using coat color

Using QTL mapping, we identified the genetic region, and ultimately strong candidate genes, mediating tolerance in the CC024 mice. Since CC024 and CC061 are fully homozygous, and their genomes are fully sequenced, we used the Mini Mouse Universal Genotyping Array (MiniMUGA) to genotype the F2 population. MiniMUGA has 11000 markers that differentiate around 120 classical inbred lines and are equally spaced across the genome, enabling precise QTL mapping (Sigmon et al., 2020). We used the coat color phenotype of our F2 population as a control for genotyping and QTL codes.

Our F2 population had two different coat colors: agouti and albino. These coat colors followed the Mendelian inheritance pattern of 3:1; 77% were agouti, and 23% were albino in our F2 population. We binary-coded (0 = agouti, and 1 = albino) the coat color data for our QTL analysis and identified a significant QTL peak on chromosome 7 (Fig S2A). The peak marker was located at 87.12 MB with a 1.5 LOD drop range between 83 and 89 MB. CC061 allele and the heterozygote allele caused the agouti coat color, while the CC024 allele was responsible for the albino coat color (Fig S2B). Among the eight founders, A/J contributed the CC024 allele, and C57Bl6/J contributed the CC061 allele in this region (Fig S2C).

We looked for variants in the founder alleles in this region with predicted deleterious effects on protein structure and function. We found a missense variant (rs31191169) in the A/J version of the Tyrosinase (*Tyr*) gene, changing amino acid 103 from cysteine to serine. The Sorting Intolerant from Tolerant (SIFT) score for this mutation is zero, suggesting this mutation strongly affects the Tyrosinase protein structure and function. Variants in the *Tyr* gene have previously been shown to influence coat color pigmentation in mice, including the CC (Beermann et al., 1990; Ram, Mehta, Balmer, Gatti, & Morahan, 2014). The shortlisting of the *Tyr* gene variant using coat color is quality control for our other genotyping and QTL analyses.

### Genes on chromosome 7 control survival and weight change after infection in CC024 mice

Survival, weight change, and colonization were considered complex traits for QTL analysis using F2 mice. A significant QTL peak on chromosome 7 was linked to survival (Fig 5A). We named this QTL peak 24SMI (CC024 Survival after MRSA Infection). The 1.5 drop confidence interval was between 4 and 25 MB, with a peak at 15.91 MB. As expected, the CC024 allele increased survival, while the CC061 and the heterozygous allele (CC061/CC024) reduced survival after infection (Fig 5B and 5C). In the confidence interval region, the WSB allele contributed to tolerance in CC024 mice, and the C57Bl/6 allele contributed to the sensitivity phenotype of CC061 mice (Fig 5B). We identified a significant peak on chromosome 7 linked to weight change after infection (Fig S3A). We called this peak 24WMI (CC024 Weight change after MRSA Infection). The peak was located at 22.79 MB, with a confidence interval between 13 and 25 MB. This weight peak completely encompassed the survival peak and had the same founder effect pattern (Fig S3B and S3C). This result was not surprising as survival and weight change are very highly correlated (R = 0.67) (Fig S1A).

**Fig 5.**
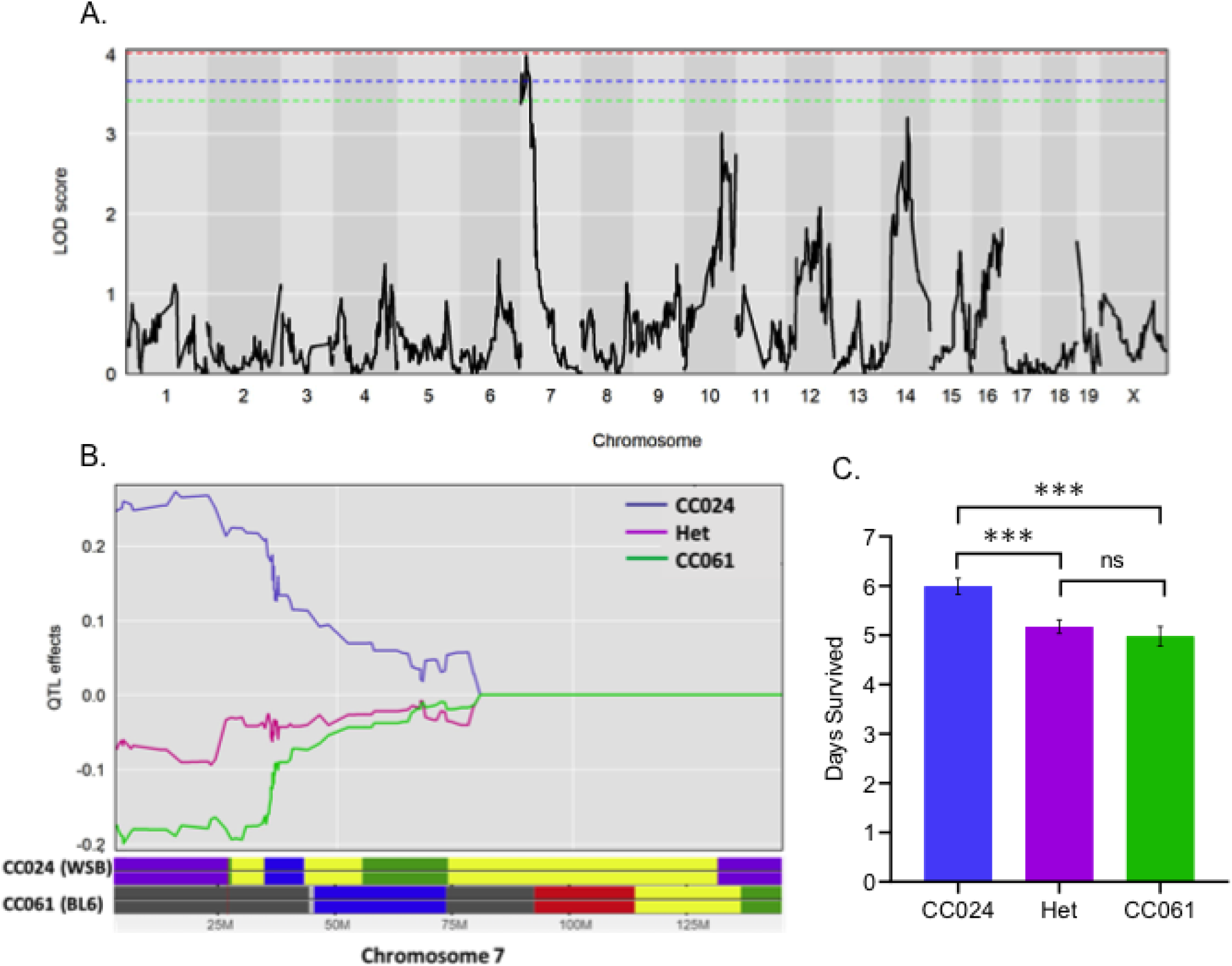
Survival after infection has a significant peak on chromosome 7. A. LOD plot for rank-transformed survival after infection (number of days). The dotted (Red – 95%, Blue – 90%, Green – 85%) lines represent the significant LOD scores for 999 permutations. B. Founder allele plot for chromosome 7. C. Founder allele contribution at the highest marker on chromosome 7, mean and standard error of the mean are shown. Het – Heterozygous allele for CC061 and CC024. Student t-test was performed (ns – no significance, *** = P < 0.001)

### Prioritizing variants on chromosome 7

We used the Mouse Phenome Database (MPD) to prioritize variants (SNPs, insertions, or deletions) within our confidence interval linked to survival and weight loss. This 21 MB region has ∼35,000 variants between the WSB and C57BL/6 alleles. We used Variant Effect Predictor (VEP) to shortlist variants within genes that were predicted to significantly impact protein structure and function. We identified 12 genes in this region with predicted high-impact variation on the structure and/or function of the predicted protein (Table S3). The variant with the highest impact for each gene and previous immune system process associations in the Mouse Genome Informatics (MGI) database (highlighted in red) are shown in Figure 6A (Fig 6A). Using the total RNA sequencing data from the kidney of F2 mice, we further prioritized these high impact variants.

**Fig 6.**
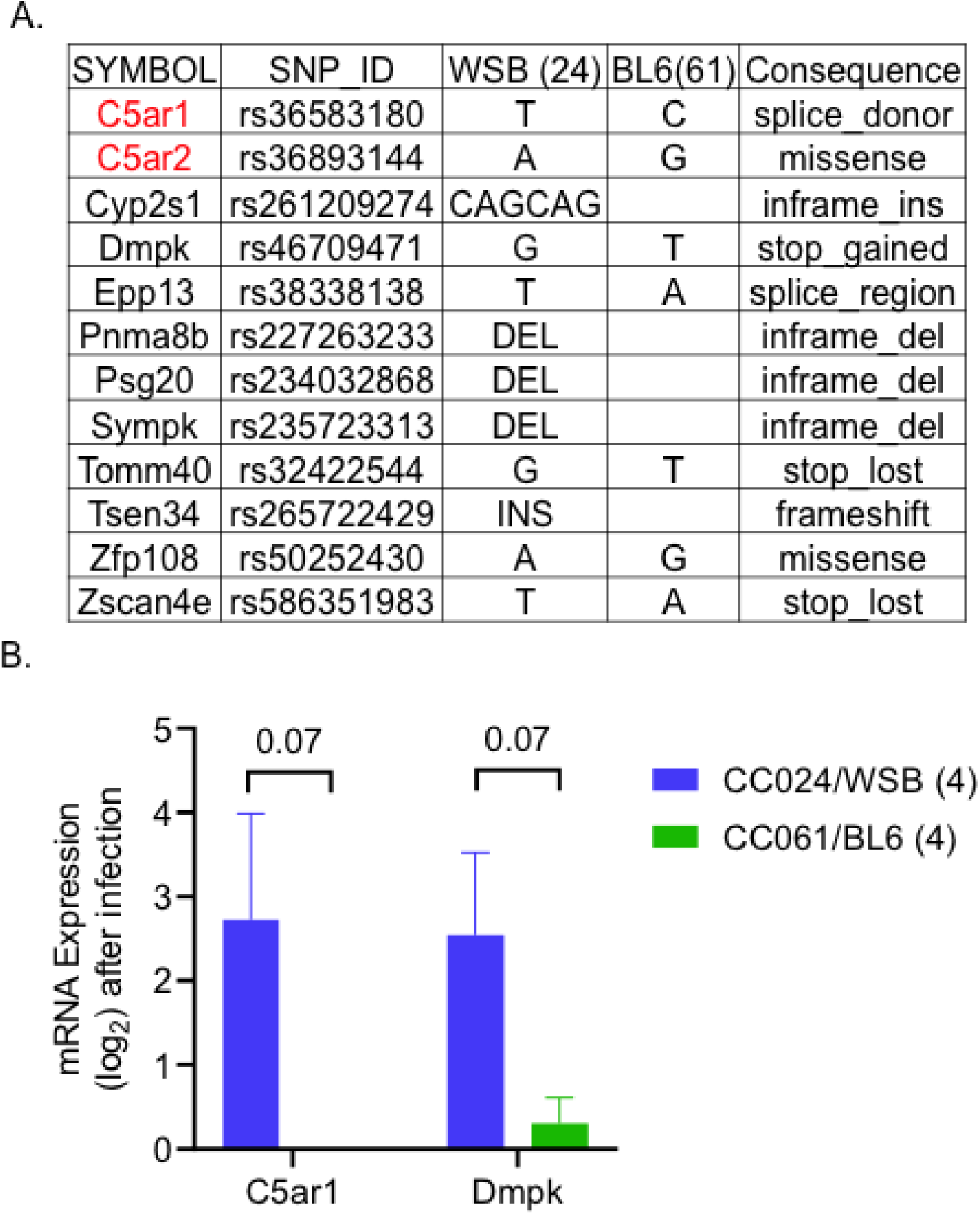
Prioritizing genes on chromosome 7 for survival after infection. A. Genes with high-impact variants shortlisted using the founder effect pattern. B. Kidney mRNA expression values after infection (log-transformed). Mean and standard error are shown. The number of animals is given in brackets. Student’s t-test was performed (Mean with standard error and P values are shown)

The mRNA expression levels of these 12 genes were not significantly different between F2 animals with a WSB and C57BL/6 allele at the peak location. This finding may be due to a low number of samples (four mice per group), or these variants may not affect transcript abundance. The closest variants to significance were complement component 5a receptor 1 (*C5ar1*) and dystrophia myotonica-protein kinase (Dmpk) genes, with P-values of 0.07. Both genes were expressed more highly in F2 animals having WSB allele compared to the C57BL/6 allele at the QTL peak (Fig 6B). Among the shortlisted genes, both *C5ar1* and *C5ar2* are essential genes in the complement cascade and are involved in immune response to infections (Yan & Gao, 2012). None of the other candidate genes have any previously associated immune system function in the MGI database. Thus, based on previous associations with the immune system, enrichment of complement cascade in CC024 mice and F2 kidney mRNA expression data, *C5ar1* and *C5ar2* are our top candidate genes for involvement in the tolerance phenotype in the 24SMI (survival) QTL peak. The *C5ar1* gene in CC024 mice has a splice donor variant that can disrupt RNA splicing resulting in an altered protein sequence and structure. Similarly, *C5ar2* in CC024 mice has a predicted high-impact missense variant with a SIFT score of 0.03. The modified version of *C5ar1* and *C5ar2* in CC024 mice contributed by the wild-derived WSB founder may provide an added advantage to survival relative to the C57Bl6 allele present in CC061 mice.

## Discussion

Host genetics plays a major role in protection against bacterial infections (Kwok, Mentzer, & Knight, 2021). Infection outcome in mice is also heavily dependent on the strain of inbred laboratory mice used, suggesting strong influence of host genetics (von Kockritz-Blickwede et al., 2008). Furthermore, mice lacking Nod2 (Nucleotide-binding oligomerization domain-containing protein 2), Myd88 (Myeloid differentiation primary response protein), and Tlr2 (Toll-like receptor 2) are more susceptible to infection with *S. aureus* than mice expressing these genes (Deshmukh et al., 2009; Takeuchi, Hoshino, & Akira, 2000). We investigated the genetic factors behind the tolerance to infections with MRSA USA300. We chose a susceptible strain (C0061) and a tolerant strain (CC024) from our previous MRSA infection screen to identify genomic regions that are involved in these phenotypes (Submitted: mBio01812-23). The tolerant strain, CC024, survived longer and lost less weight than C0061, despite similar colonization including high colonization in the kidney.

After accounting for baseline strain differences, several serum cytokines differed between CC061 and CC024 mice post infection (Fig 2C). IL-10 is the only cytokine that was increased in the serum of CC061 mice. IL-10 is produced by both innate and adaptive immune cells and is a major immunoregulatory cytokine essential for modulating the production of several other cytokines (Kühn, Löhler, Rennick, Rajewsky, & Müller, 1993; Leech, Lacey, Mulcahy, Medina, & McLoughlin, 2017; Sabat et al., 2010). Overexpression of IL-10 worsens the outcome of bacterial infections, including *S. aureus* infections (Leech et al., 2017). IL-10 creates an anti-inflammatory environment by inhibiting the production of several pro-inflammatory cytokines and chemokines (Moore, de Waal Malefyt, Coffman, & O’Garra, 2001). IL-10 inhibits T-helper cell responses (Frodermann et al., 2011), and down regulates MHC class II expression (Sendide et al., 2005). IL-10 also down regulates co-stimulatory molecules on antigen-presenting cells (Buelens et al., 1995), further decreasing T-cell activation (Z. Li, Peres, Damian, & Madrenas, 2015). The increased production of IL-10 by CC061 mice may create an anti-inflammatory state that contributes to reduced survival of these mice during MRSA infection. Whether *S. aureus* itself manipulates IL-10 levels to evade the innate immune system is not clearly understood. Elevated IL-10 levels likely contribute to the severity of MRSA infection in CC061 mice.

Eight cytokines were increased after infection in serum of MRSA tolerant CC024 mice, after accounting for pre-infection baseline differences. Pro-inflammatory cytokines IL-17A and IL-22 were increased in the serum of tolerant CC024 mice after MRSA infection. IL-17A recruits neutrophils to the site of infection (Ye et al., 2001). Though IL-22 belongs to the IL-10 family, it plays a major role in controlling *S. aureus* nasal colonization by inducing antimicrobial peptide (AMP) production (Mulcahy, Leech, Renauld, Mills, & McLoughlin, 2016). In bacterial infections, both IL-17 and IL22 promote AMP production; IL-22 mainly acts by increasing barrier function, while IL-17 aids in neutrophil recruitment (Aujla et al., 2008; Mills, 2023). Increased serum levels of these cytokines in CC024 mice likely contribute to the longer survival in these mice after MRSA infection.

Both IL-1β and IL-18, elevated in CC024 mice after infection, act on lymphoid cells and enhance the production of IL-22 (Bergmann et al., 2017; Victor et al., 2017). In addition, IL-1β also inhibits the production IL-10 by memory TH17 cells (Zielinski et al., 2012), thereby counteracting its adverse effects. In *S. aureus* skin infections, both IL-4 and IL-23 (elevated in CC024) promote the production of IL-17 (Cho et al., 2010; Leyva-Castillo et al., 2021). IL-27 is a pleiotropic cytokine that has both pro- and anti-inflammatory properties. Though IL-27, along with IL-10, has been shown to enhance nasal colonization and increase susceptibility to *S. aureus* pneumonia (Kelly, Leech, Doyle, & McLoughlin, 2022; Robinson et al., 2015), its role in suppressing inflammation in the later stages of infection may be beneficial in CC024 mice (Beizavi, Zohouri, Asadipour, & Ghaderi, 2021). Overall, decreased levels of IL-10, along with increased levels of IL-17A and IL-22, may play a role in the survival of CC024 mice after MRSA infection. In a recent *S. aureus* vaccination study, increased levels of IL-10 and inhibited IL-17 production reduced the efficiency of the vaccine (Narita, Hu, Asano, & Nakane, 2019). This study further supports important roles for IL-10 and IL-17 in disease outcome after *S. aureus* infection.

The complement and coagulation cascade pathway is up-regulated in CC024 mice, both before and after MRSA infection (Fig 3). The complement system is one of the first host defense systems that *S. aureus* encounters during infection (Wójcik-Bojek et al., 2022). Complement consists of more than 30 proteins in plasma that opsonize bacteria, depositing complement activation products on the bacterial surface and attracting immune cells (de Jong, van Kessel, & van Strijp, 2019). Complement can be activated by three pathways, which converge at the formation of *C3* convertase that cleaves *C3* into C3a and C3b (Dunkelberger & Song, 2010). C3b opsonizes the *S. aureus* surface, while C3a activates immune cells, including macrophages and T cells (Wójcik-Bojek et al., 2022). *C3* deficient mice are more susceptible to *S. aureus* infections (Cunnion, Benjamin, Hester, & Frank, 2004; Na et al., 2016). While the mechanism underlying the baseline up-regulation of the complement genes, *C3* and *C4b* in CC024 mice is not understood, these immune factors may improve CC024 survival by reducing the initial pathogen load.

To identify the genes needed for CC024 mice to tolerate MRSA infection despite high colonization in some organs, we generated an F2 cross between CC024 and CC061. Our data suggest that the genetic factors influencing MRSA colonization are well segregated in the F2 generation, and a complex trait controls their interaction. Survival in F2 mice ranged from 2 to 7 days post-infection, with a median of 6 days (Fig 4A). Kidney colonization in F2 animals was roughly equivalent, irrespective of the euthanizing day (day 2 vs. day 6, Fig S1C). Thus, F2 mice appear to be tolerant, as they can differentially survive the infection with similar bacterial loads. Survival correlated with kidney colonization (R = -0.57) and was considered a complex trait for QTL mapping.

The top candidate genes for the survival and weight change QTL in our analysis are the complement factor *C5a* receptors, *C5ar1* and *C5ar2*. In the earlier stages of infection, activation of the complement system is important to clear the pathogen (Schifferli et al., 1986). However, during sepsis and later stages of infection, over-activation of complement negatively impacts the immune system (Guo et al., 2003).

Among the complement factors, *C5a* is the most potent inflammatory mediator to increase both pro-inflammatory factors, including TNF-α, IL-1β, IFN-γ, IL-6, and IL-8 and anti-inflammatory factors, including IL-10, IL-13, IL-4 and TGF-β (Flierl et al., 2008; Guo, Riedemann, & Ward, 2004; Le Tulzo et al., 2002; Titheradge, 1999; Wolkow, 1998). Activation of the complement system leads to the cleavage of C5 into *C5a* and C5b (Horiuchi & Tsukamoto, 2016). *C5a*, which binds both *C5ar1* and *C5ar2*, is an anaphylatoxin and a chemotactic factor triggering an inflammatory response (Colley et al., 2018; Ricklin & Lambris, 2013).

In mice and humans, *C5ar1* and *C5ar2* share high sequence identity but differ significantly in important regions that negatively affect the function of *C5ar2* (X. X. Li, Lee, Kemper, & Woodruff, 2019). Conflicting evidence exists on the potential roles of *C5ar2*: is it a decoy receptor to *C5ar1* with anti-inflammatory properties, or a unique signaling receptor with pro- or anti-inflammatory properties (T. Zhang, Garstka, & Li, 2017)? *C5ar1* is expressed in both myeloid cells, including macrophages, monocytes, and neutrophils and non-myeloid cells, including Kupffer cells, astrocytes, endothelial cells, and kidney tubular epithelial cells (Bosmann, Sarma, Atefi, Zetoune, & Ward, 2012; Werfel et al., 1992) (Drouin et al., 2001; Sun et al., 2009; Zahedi et al., 2000). During sepsis, expression of *C5ar1* is up-regulated in organs, including the lung, liver, heart, and kidney (Riedemann et al., 2002).

*C5ar* deficient mice have increased survival in cecal ligation and puncture (CLP) induced sepsis (Rittirsch et al., 2008) and increased resistance to bacteremia and endotoxic shock (Hollmann, Mueller-Ortiz, Braun, & Wetsel, 2008). Similarly, a *C5ar1* antagonist increased survival in CLP-induced sepsis (Huber-Lang et al., 2002) and reduced bacterial counts in several organs (Riedemann et al., 2002). These data suggest that activation of *C5ar1* hastens the progression of sepsis. Binding of *C5a* to both *C5ar1* and *C5ar2* promotes inflammation and tissue damage (K. Li et al., 2017; Peng et al., 2019; T. Zhang et al., 2020). Binding of *C5a* to these receptors results in increased production of several cytokines, chemotaxis of immune cells, intracellular calcium release, and superoxide anion production (Postma et al., 2004; Yan & Gao, 2012) that results in a cytokine storm damaging vital organs (Fajgenbaum & June, 2020).

In CC024 mice, *C5ar1* and *C5ar2* have mutations that are predicted to significantly alter the structure of these *C5a* receptor proteins (Fig 6A). *C5a* may be either unable to or bind differentially to, the altered *C5ar1* and *C5ar2* receptors in CC024 mice, reducing inflammation, reducing the cytokine storm, reducing the resulting tissue damage, and potentially leading to tolerance. This hypothesis is also consistent with our finding that heterozygotes at these loci survive poorly (Fig 5B and 5C); a single CC061 allele for one of these receptors may be sufficient to allow activation of the complement system, leading to reduced survival in heterozygotes. These *C5ar1* and *C5ar2* variants, independently or together, are strong candidates for an important role in the increased survival during MRSA infection of CC024 mice.

Successfully eliminating a pathogen while minimizing damage to host tissue requires timing and coordination of the host immune system. Pathogens have evolved to take advantage of imperfections in the immune system. Here, natural variants occurring within the complement system could benefit the host by reducing inflammation and tissue damage in certain situations, thereby increasing the ability of the host to tolerate an infection for longer periods of time. With growing interest in targeting the complement system for developing therapeutics against sepsis (Garred, Tenner, & Mollnes, 2021; Rodrigues, Picco, Morgan, & Ghazal, 2021; Sommerfeld et al., 2021), how natural variants of the complement system influence the effects of this system may be a fruitful avenue of future study.

## Materials and Methods

### Ethics statement

All mouse studies followed the Guide for the Care and Use of Laboratory of Animals of the National Institutes of Health (National Research Council Committee for the Update of the Guide for the & Use of Laboratory, 2011). The animal protocols (2015-0315 D and 2018-0488 D, 2019-0411) were reviewed and approved by Texas A&M Institutional Animal Care and Use Committee (IACUC).

### Bacterial strains and media

Methicillin-resistant *Staphylococcus aureus* isolate (MRSA USA300) used in this study was the kind gift of Dr. Magnus Hook (Texas A&M Institute of Biosciences and Technology, Houston). USA300 is a fully virulent, community-acquired clone of MRSA. Strains were routinely cultured in Luria-Bertani (LB) broth and plates supplemented with antibiotics when needed at 50 mg/L Kanamycin Sulphate. For murine infections, strains were grown aerobically at 37°C to a stationary phase in LB broth supplemented with kanamycin.

### Murine strains – breeding and genotyping

Collaborative Cross (CC) mouse strains – CC061 and CC024, F1s (CC061xCC024), and F2s (CC061xCC024) were used for these experiments (Table S1). The CC strains were initially purchased from UNC’s Systems Genetics Core Facility (SGCF) and were bred at the Division of Comparative Medicine at Texas A&M University to generate F1 and F2 generation (Table S1). Mice were bred in trios (2 females and one male), and offspring were weaned at 21 days. Tail snips were collected during weaning and genotyped using the Mini MUGA panel at Neogen (Sigmon et al., 2020). Mice were fed Envigo Teklad Global 19% Protein Extrudent Rodent Diet (2919) and were provided with cardboard huts, food, and water ad libitum.

### Parental strain data

Data for the two parental CC strains, CC024 and CC061, were collected from another 1-week experiment performed previously (Submitted: mBio01812-23). Data from our earlier study includes survival after infection, organ colonization, complete blood count (CBC), and cytokine/chemokine levels (Table S2), as the basis of the study and for comparison with F1 and F2 generation data. Samples for this analysis were obtained as previously described (Submitted: mBio01812-23).

### Health monitoring

For F1 and F2 mice, we manually scored five health parameters twice daily: weight, physical appearance, body conditioning, and provoked and unprovoked behavior. The scoring scale ranged from 0-3, with zero = normal and three = abnormal. Additionally, as a measure of activity, four small nestlets were placed at each corner of the cage in the evening. The following day, the number of nestlets moved from the cage corners into the hut was recorded.

### MRSA Infection

After baseline scoring for health, F1 and F2 mice were injected with MRSA USA300. Briefly, Mice were anesthetized using isoflurane and injected with 1 x 10^7^ CFU in 50 µl of LB broth intravenously at the inferior fornix into the retro-orbital sinus. Mice that became moribund within 6 hours of infection were humanely euthanized and removed from the experiment.

### Bacterial load determination

After infection, mice were monitored twice daily using visual health scoring. Mice that developed illness as measured by the health score were humanely euthanized by CO_2_ asphyxiation. After euthanasia, kidneys were collected. A consistent region of the right kidney was collected in 3mL ice-cold PBS and homogenized. The serially diluted homogenate was plated on LB plates supplemented with kanamycin for bacterial enumeration. Data are expressed as CFU/g of tissue.

### QTL analysis

Data from 315 F2 mice were included in the analysis. QTL analysis was performed using R/qtl2 software (K. W. Broman et al., 2019). This method accounts for the complex population structure in CC strains. Briefly, genome scans were performed on the rank-transformed phenotype. The generated Logarithm of Odds (LOD) score is the likelihood ratio comparing the hypothesis of a QTL at a given position versus that of no QTL. Each phenotype was randomly shuffled 999 times to establish genome-wide significance, and LOD scores were calculated for each iteration. The 85th percentile of the scores was considered significant for that phenotype. The genomic confidence interval was calculated by dropping the LOD scores by 1.5 for each significant peak. Mouse Genome Informatics (MGI) was used to find the genes and QTL features within each interval (Blake et al., 2021; Bogue et al., 2023). To further shortlist candidate genes, the founder strain distribution pattern was queried against the CC variant database (V3 Version) (Karl W Broman, 2017). The Variant effect predictor (VEP) from the ensemble was used to calculate the impact score for the variant (Cunningham et al., 2022).

### RNA extraction and sequencing

Total RNA was extracted from frozen tissues using Direct-zol RNA Miniprep plus kit following the manufacturer’s protocol (Zymo Research - R2073). The purity and quantity of the extracted RNA were analyzed using RNA 6000 Nano LabChip Kit and Bioanalyzer 2100 (Agilent CA, USA, 5067-1511). High-Quality RNA samples with RIN number > 7.0 were used to construct the sequencing library. Following the vendor’s recommended protocol, we performed the 2×150bp paired-end sequencing (PE150) on an Illumina Novaseq™ 6000.

### RNA seq data analysis

Reads were trimmed using trim galore (version 0.6.7) (Krueger, 2019). This procedure removed adapters, poly tails, more than 5% of unknown nucleotides, and low quality reads containing more than 20% of low-quality bases (Q-value <20). Both forward and reverse hard trimmed at 100 base pairs. FastQC was used to verify data quality before and after cleaning (Andrews, 2010; “FastQC,” 2015). Cleaned reads were aligned and counted against the mouse reference genome (GRCm39) using STAR (version 2.7.9a) aligner (Dobin et al., 2013). Downstream processing of the data was performed using IDEP 1.0 (S. X. Ge, Son, & Yao, 2018; X. Ge, 2021). Gene counts were analyzed for differentially expressed genes using DESeq2 (Love, Huber, & Anders, 2014).

### Data availability

All of the raw data associated with each mice used in the supplementary materials (Table S1 and S2). RNA sequencing is available in Gene Expression Omnibus (currently being uploaded ####).

## Acknowledgements

We thank Dr. Magnus Hook for providing the bacterial strain used in this study. We also would like to thank Dr. Joana Rocha, Mr. Connor Mathis, and Mrs. Kaya Mariello for technical assistance. This work was funded by the Defense Advanced Research Projects Agency grant number D17AP00004.

---

**Fig S1. Infection outcome in F2 cross.** A. Heat map showing Spearman correlation ‘R’ values between survival, weight change, Infection dose, and kidney CFU. B. Survival after infection separated by males and females. C. Kidney colonization of F2 mice by days survive. The median and interquartile range are shown. B – Student’s t-test and C – Mann-Whitney tests were performed (ns-no significance, *** = P < 0.001).

**Fig S2. QTL for coat color in the F2 population.** A. LOD plot for coat color after infection (0 – Agouti, 1 – Albino/white). The dotted (Red – 95%, Blue – 90%, Green – 85%) lines represent the significant LOD scores for 999 permutations. B. Founder allele plot for chromosome 7. C. Founder allele contribution for chromosome 7.

**Fig S3. QTL for weight change after infection in the F2 population.** A. LOD plot for rank-transformed weight change after infection. The dotted (Red – 95%, Blue – 90%, Green – 85%) lines represent the significant LOD scores for 999 permutations. B. Founder allele plot for chromosome 7. C. Founder allele contribution for chromosome 7.

Table S1: Mouse strains used in this study. Number of mice, origin, and location of breeding for all the mice used in the study.

Table S2: Data associated with each mouse. All the pre-infection and post-infection data were associated with each mouse used in this study.

Table S3: Shortlisted variants for QTL peak on chromosome 7 for survival after infection. Table S4: Enriched KEGG pathway genes

